# Signatures in CRISPR Mutational Spectra Reveal Role and Interplay of Genes in DNA Repair

**DOI:** 10.1101/2025.11.25.690381

**Authors:** Colm Seale, Marco Barazas, Robin van Schendel, Marcel Tijsterman, Joana P. Gonçalves

## Abstract

**Motivation:** Understanding double-strand break (DSB) repair and its disruption is key to decipher genomic instability driving diseases such as cancer and reveal therapeutic avenues. Numerous genes have been linked to DSB repair for the first time in recent genome-wide perturbation studies assessing effects on mutational outcomes. However, the functional roles of most such genes remain poorly understood. Evidence from other studies shows that related genes similarly modulate the frequency of specific mutational outcomes following DSB repair, but analysis has largely ignored the multiplicity of gene functions and cross-talk between pathways. Here, we infer functional roles for candidate genes based on knockout mutational spectra by identifying mutational signatures shared with known genes and then mapping them to DSB repair functions. Signatures are identified using non-negative matrix factorization (NMF) to reflect functional multiplicity and cross-talk.

**Results:** We generated mutational spectra for mouse embryonic stem (mES) cells at three target sites across CRISPR knockouts of 307 known and 459 candidate DSB repair genes. Analysis using NMF revealed four mutational signatures associated with homology-directed repair (HDR), microhomology-mediated end-joining (MMEJ), and the initiation and ligation components of non-homologous end-joining (NHEJ). Signatures suggested that candidate genes *Dbr1* and *Hnrnpk* could influence MMEJ and Fanconi anaemia (FA), and that loss of core NHEJ components (e.g. MRN complex or Ku proteins) could shift repair preference towards Ku-*independent* NHEJ. These findings demonstrate the utility of NMF for characterizing the contribution of genes to repair pathways and provide a foundation to discover new gene functions in DSB repair.

**Availability:** github.com/joanagoncalveslab/MuSpectraNMF.

## Introduction

Effective cellular repair of DNA double-strand breaks (DSBs) is essential to prevent genomic instability leading to cell death or the development of diseases such as cancer (Cannan and Pederson, 2016). Deficiencies in DNA damage response do not only drive tumor progression, but also expose vulnerabilities of tumor cells that can be leveraged for treatment. Conventional chemo and radiation therapies work by inducing substantial DNA damage, from which tumor cells with compromised repair struggle to recover. However, such therapies also affect healthy cells, causing debilitating side effects and possibly recurrence. Targeted treatments mitigate this toxicity by design by selectively exploiting unique handicaps of tumor cells. For instance, PARP inhibitors trigger the joint essentiality between PARP and BRCA genes to treat tumors with deficient BRCA function (Chen, 2011). While the benefit of targeted therapies can be great, development requires in-depth knowledge of DSB repair mechanisms and the functions of individual genes. Advancing the understanding of DSB repair is therefore essential to reveal mechanistic relationships offering new therapeutic possibilities to improve patient care and quality of life (Awwad *et al*., 2023).

The study of DSB repair has been accelerated by the availability of CRISPR (Clustered Regularly Interspaced Short Palindromic Repeats) technology to induce DSBs at predefined genomic loci (Jinek *et al*., 2012). A growing number of studies combine CRISPR targeting with DNA sequencing readout to analyze mutational spectra resulting from DSB repair activity and gain new insights into repair mechanisms at scale. Specifically, several studies have used CRISPR targeting across collections of loci to investigate the influence of sequence context around the DSB within a few different cell types or genomic contexts (van Overbeek *et al*., 2016; Shen *et al*., 2018; Shou *et al*., 2018). Others have paired systematic knockouts of individual DSB repair genes with CRISPR targeting across loci to tease out the function of each gene and its specific contribution to DSB repair mechanisms (Hussmann *et al*., 2021). For example, cells harboring knockouts of *Polq* show significantly fewer long deletions with microhomologies (MHs) at DSBs post-repair than wild-type cells, consistent with the key role of the gene in the microhomology-mediated end-joining repair pathway (Wyatt *et al*., 2016; Hussmann *et al*., 2021). Such insights have also recently inspired genome-wide and follow-up focused screens of gene knockout effects on mutational spectra aimed at uncovering new gene associations with DSB repair (Barazas *et al*., 2025; Seale *et al*., 2025).

Existing studies to elucidate the role of genes in DSB repair take different approaches to analyze the effects of the gene knockouts on mutational spectra relative to controls. One strategy is to separately examine the impact on individual outcomes of interest, such as 1bp insertions or deletions (Barazas *et al*., 2025). Such univariate analysis ignores covariation of outcomes produced by shared underlying mutational processes, whereas repair mechanisms and genes do not function in isolation. Multivariate methods are also applied to reveal genes involved in DSB repair through outlier analysis of spectra from genome-wide knockout screens, resulting in candidate associations of genes with DSB repair but no further insight into possible functions (Seale *et al*., 2025). Alternative multivariate strategies are further used to discover relationships between known DSB repair genes based on similarity of their mutational spectra, relying on conventional clustering or manifold learning. Conventional clustering techniques such as hierarchical clustering (Ran *et al*., 2023) focus on global similarities across the entire mutational spectra and thus ignore that the same gene or mechanism can contribute to multiple outcomes, whereas manifold learning methods like UMAP (McInnes *et al*., 2018) can capture local relationships between genes based on subsets of outcomes but are not straightforward to interpret due to their non-linear nature (Pal and Sharma, 2020; Kobak and Linderman, 2021).

Here, we propose using non-negative matrix factorization (NMF) to analyze mutational spectra of known and candidate DSB repair genes, and identify relationships offering insight into the functional role of such candidates. The advantage of NMF is that it captures local patterns while being interpretable and producing clusters or factors with “soft” gene memberships, reflecting the fact that genes can operate in multiple DSB repair pathways and that distinct pathways can produce identical outcomes at varying rates (also co-occurring with different sets of other outcomes). This is achieved by decomposing the mutational spectra of all gene knockouts into mutational signatures capturing patterns across subsets of knockouts, and signature exposures quantifying the contribution of each signature to the spectrum of each gene knockout (Lee and Seung, 1999; Koh *et al*., 2021).

In this work, we generate mutational spectra for three target sites under individual knockouts of 766 genes, including 307 known DSB repair genes and 459 candidates selected based on outlying spectra from a previous genome-wide knockout screen (Seale *et al*., 2025). We investigate the ability of NMF to identify signatures and exposures that recover known mutational patterns and responsible genes linking signatures to DSB repair pathways. We also leverage signature depletions to suggest DSB repair mechanisms for candidate genes, and explore how joint depletion patterns may provide further functional granularity for genes of the same pathway (Fig. 1).

**Fig. 1:**
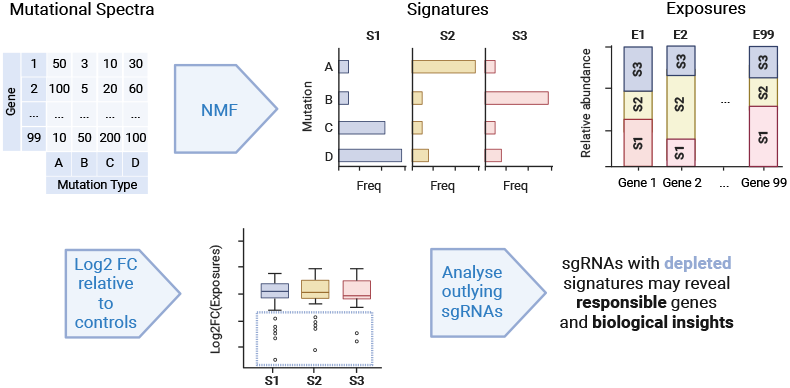
Identifying DSB repair gene function via NMF analysis of mutational spectra. Mutational spectra, defined as counts over a set of unique mutational outcomes (columns A–D), for cell populations carrying knockouts of individual genes (rows E1-E99). Estimation of mutational signatures (S1-S3) and associated exposures (E1-E99) using NMF. Analysis of signature exposure ratio relative to controls (log2 fold-change) identifies gene knockouts depleted in signature-specific mutational outcomes, which can be further linked to other genes and functional mechanisms with impact on the same signatures.

## Methods

We analyzed two datasets of mutational spectra for mouse embryonic stem (mES) cells carrying knockouts of individual genes: the primary dataset, generated for this study, comprising knockouts of 766 genes and three target sites; and a published dataset, which we name Barazas2025 (GEO accession: *unreleased*), comprising knockouts of 742 genes and one target site Barazas *et al*. (2025). There were 273 genes in common between the two screens. We included the Barazas2025 dataset mainly to assess the robustness of the mutational signatures identified from the primary dataset.

### Generating mutational spectra

#### Selecting genes for the primary screen

The set of 766 genes for the primary screen consisted of 459 candidate genes and 307 known DSB repair genes. The candidate genes were based on the analysis of a genome-wide study quantifying the effect of individual gene knockouts on mutational spectra (Seale *et al*., 2025). While the genome-wide study traded gene coverage for mutational spectra resolution, here we focused on profiling the more promising genes at higher resolution to increase the robustness of potential findings. Specifically, we selected the 459 genes with the largest impact on their knockout mutational spectra according to a multivariate outlier score measuring deviation relative to the center of the distribution of all mutational spectra. To be able to relate mutational signatures and candidate genes to established DSB repair mechanisms, we additionally selected 307 DSB repair genes as the union of the following three sets of genes: *(i)* annotated with Gene Ontology (GO) terms “double-strand break repair” or “interstrand cross-link repair” (Consortium, 2021); *(ii)* annotated with KEGG pathways “Homologous recombination” (ID: mmu03440), “Non-homologous end joining” (ID: mmu03450), or “Fanconi anaemia pathway - Mus musculus (house mouse)” (ID: mmu03460) (Kanehisa *et al*., 2023); and *(iii)* curated as the 118 DSB repair genes with the largest effect in the Repair-seq study (Hussmann *et al*., 2021).

#### Screening DSB repair outcomes

To generate mutational spectra, we followed the experimental protocol in Barazas *et al*. (2025) (Supplementary Fig. S1). Briefly, paired guide-target DNA sequences were integrated into the genomes of endogenously Cas9-expressing mouse embryonic stem (mES) cells via lentiviral transduction. Each integrated sequence contained a single-guide RNA (sgRNA) designed to knock out a specific gene or as a non-targeting control, and one of three 20bp target sequences with protospacer adjacent motif (denoted T1-T3). For the primary screen, we used a total of 2,321 knockout sgRNAs (~3 per gene), and 50 additional non-targeting control sgRNAs. The Barazas2025 screen included a total of 2,400 knockout sgRNAs for 742 genes, and 170 non-targeting control sgRNAs. After allowing 5 days for genomic integration and gene knockout, we split the bulk cell population into nine samples for the primary screen (3 targets × 3 replicates) or 3 samples for the Barazas2025 screen (1 target, 3 replicates). Per sample, we performed a second round of lentiviral transduction to express sgRNAs and induce DSBs at the integrated target sites. Following ~80h of cell culture for cleavage and repair, we used paired-end DNA sequencing to characterize repair outcomes at the integrated sites.

#### Processing outcome sequences into mutational spectra

We used the SIQ v4.3 tool (van Schendel *et al*., 2022) to call mutational outcomes from the sequencing data, with parameters “-m 2 -c -e 0.05” (requiring a minimum number of 2 reads to count an event, collapsing identical events to a single record with the sum of counts, and allowing a maximum base error rate of 0.05). The outcomes were split into four categories: deletion; insertion; templated insertion, denoting a deletion with an insertion where the inserted sequence matches a region flanking the cut site; and HDR event, for any insertion matching a provided donor template DNA. Additionally, we recorded length and location for both insertions and deletions, as well as microhomology (MH) length when present for deletions. All templated insertions were collapsed into a single category, with the respective sum of counts over all observed events. Wild-type sequences were excluded to produce a final collection of mutational outcomes with the corresponding counts per sgRNA.

To filter rare outcomes, we excluded outcomes with a geometric mean frequency below 0.002 across the non-targeting controls. This resulted in a final set of mutational spectra containing 28-44 outcomes per target site and replicate for the primary screen and 32 outcomes per replicate for the Barazas2025 screen (Supplementary Table S1 for outcome counts, and Supplementary Tables S2-S5 for mutation details).

#### Replicate quality analysis and selection

We assessed data quality by calculating pairwise Pearson’s correlations between replicates of the same target site, using the sgRNAs and outcomes common to each pair. Average correlations were above 0.98, indicating high quality throughout (Supplementary Table S5). We selected the replicate with the largest number of recovered gene knockouts per target site (T1: 2,291, T2: 2,291, T3: 2,304, Barazas2025: 2,358; Supplementary Table S1) for downstream analysis.

### Identifying co-occurring mutational patterns

#### Analyzing patterns across target sites

Since mutational spectra are target site-specific, to characterize patterns across sites for the primary dataset, we concatenated the mutational spectra of the three target sites per sgRNA into a single mutational spectrum covering all three sites. We excluded sgRNAs missing from any of the target sites. To mitigate target-specific batch effects, we scaled the counts per sgRNA to equalize the ranges of counts across the different targets. This resulted in a three-target mutational spectra count matrix of 2,288 sgRNAs × 112 mutational outcomes.

#### Identifying signatures and exposures

Each mutational spectrum reflects the aggregated contribution of all mutational processes, including DSB repair mechanisms, active in the cell population from which the spectrum was derived. Each mutational process tends to produce specific mutational outcomes at specific rates, resulting in a fingerprint or signature. Our goal is to decipher the mixture of signatures underlying the collection of mutational spectra using NMF, and later attempt to link each signature to responsible genes and pathways.

Formally, the input is a matrix *V ∈* ℕ^*m×n*^ of *n* mutational spectra defined as count distributions over a set *M* of *m* mutational outcomes. We aim to decompose *V* into the product of two matrices *V* ≈ *S × E*: the signature matrix *S ∈* ℝ^*m×k*^, representing a collection of *k* signatures, each defined as a frequency distribution over the set of outcomes *M*; and the exposure matrix *E ∈* ℕ^*k×n*^ denoting the contribution of each of the *k* signatures to every one of the *n* mutational spectra.

Signatures *S* and exposures *E* can be estimated *V*, given a fixed number of signatures *k*, using NMF (Lee and Seung, 1999). Here, we used the SigProfilerExtractor v1.1.23 framework, which includes additional bootstrapping steps to identify more robust or stable signatures for a fixed *k*, and an evaluation procedure to optimize the number of signatures *k*. We optimized the number of signatures between a minimum of 1 and a maximum of 10, and used 30 bootstraps for signature robustness or stability.

SigProfilerExtractor selects the solution *W* and *H* for the number of signatures *k* that produces the closest reconstruction of the original mutational spectra *V*, with an average stability across bootstraps above a threshold (we used 0.8). We identified 4 signatures for the primary dataset and 5 for the Barazas2025 dataset (Supplementary Figures S1 and S2). SigProfilerExtractor assigns a string identifier to each signature, such as ‘CH112A’, where ‘CH’ indicates the signature is extracted over a set of custom outcomes, ‘112’ is the number of outcomes in the set, and ‘A’ is a letter of the alphabet identifying a specific signature.

### Elucidating responsible genes and pathways

#### Identifying genes responsible for signatures

Given that exposures indicate the contribution of the signatures to the mutational spectra, changes in exposures can be used to detect genes with an effect on those signatures. If a gene is responsible for a signature, knocking it out should weaken the strength of that pattern and therefore lower the signature exposure compared to control cells, which we term as signature depletion. To detect this type of effect, we calculate the change in exposure *δ*_*t,s*_ for a gene knockout spectrum *t* and signature *s* as:

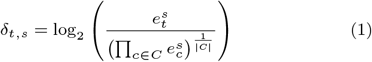

Where 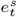 and 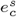 denote exposures of signature *s* for gene knockout spectrum *t* and non-targeting control spectrum *c*, respectively, and *C* refers to the set of all non-targeting control spectra of size |*C*|. Note that the denominator corresponds to the geometric mean of the exposures of the non-targeting controls. For the log2 fold change calculations, we handled zeros by applying multiplicative replacement, using defaults as implemented in scikit-bio v0.5.9 (Rideout *et al*., 2024).

#### Linking signatures to DSB repair functions

To associate signatures with biological functions, we performed GO enrichment analysis. First, we quantified the effect of each gene knockout *t* on the exposures of the different signatures relative to the non-targeting controls using the change in exposure formula *δ*_*t,s*_ (Equation 1). Second, we singled out genes “responsible” for a given signature as those genes whose knockout spectra showed outlying low change in exposure for that signature (below Q1 - 1.5×IQR, where Q1 is the first quartile and IQR is the inter-quartile range). Finally, we performed enrichment analysis for the selected “responsible” genes against the full gene set as the background, using the *GOEnrichmentStudyNS* function in “goatools” v1.2.3 (Klopfenstein *et al*., 2018). We corrected the resulting *p*-values for multiple testing using the Benjamini-Hochberg method (Benjamini and Hochberg, 1995).

#### Visualizing pathway-signature activity

To visualize the influence of gene-pathway members on exposure profiles, we obtained labels from public databases (Baldarelli *et al*., 2024; Milacic *et al*., 2024) and lists curated by major studies (Wood *et al*., 2001; Knijnenburg *et al*., 2018; Olivieri *et al*., 2020). We first compiled gene sets involved in HDR, NHEJ, and Fanconi anaemia (FA) per source. Then we used majority voting to assign one single label per gene, excluding ties to reduce ambiguity.

## Results and Discussion

### NMF identifies mutational processes and shared outcomes

Our primary aim with this study was to uncover new functional roles for genes in DNA repair by leveraging their similarity to known DSB repair genes, based on the effect of gene knockouts on mutational patterns. To assess the potential of this approach, we first investigated if we could recover associations between genes with previously validated DSB repair functions. In addition, we sought to characterize relationships between mutational outcomes and describe how related genes might share responsibility for such outcomes via repair pathway co-membership or upstream co-regulation. We analyzed global associations between genes or outcomes using hierarchical clustering and also identified more localized co-occurring mutational patterns involving subsets of genes and mutational outcomes using NMF.

Hierarchical clustering of the gene knockouts based on changes in the frequency of mutational outcomes relative to controls across the three target sites revealed functionally related groups (Fig. 2, central heatmap), including; MRN complex genes *Mre11a*/*Nbn*/*Rad50* involved in DNA damage sensing and repair, with influence on pathway choice; Lig4 complex genes involved in DNA ligation *Lig4* /*Xrcc4* /*Nhej1* /*Poll*, NHEJ-initiating Ku complex genes *Xrcc5* /*Xrcc6*; RING-type E3 ubiquitin ligases *Rnf8* /*Rnf168* involved in response to DNA damage and recruitment of repair factors with influence on pathway choice; and Fanconi anemia (FA) core complex genes. These associations confirmed that knockouts of genes with related roles tended to cluster together based on changes in the mutational spectra.

**Fig. 2:**
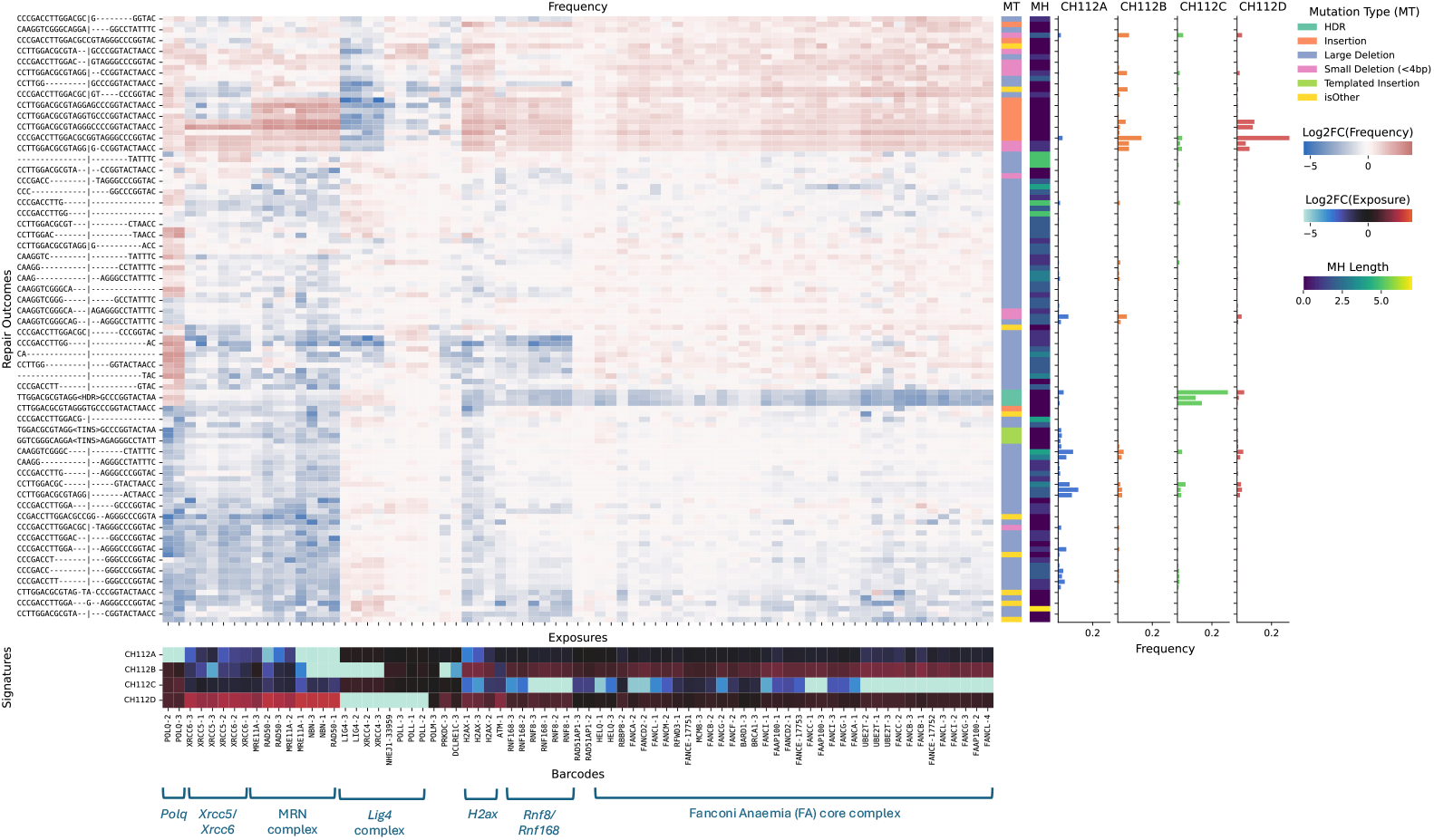
Impact of gene knockouts on mutational spectra and identified mutational signatures. (Central heatmap) Change in frequency of mutational outcomes (rows) for the 75 gene knockouts (columns) with the largest outcome redistribution compared to non-targeting controls (chi-squared-like statistic from Hussmann *et al*. (2021)). We show mutational outcomes across three target sites (rows), further characterized by mutation type (MT) and microhomology length (MH). Change in frequency quantified using the log2 fold-change in outcome frequency between the gene knockout spectrum and the geometric mean of the non-targeting control spectra. Gene knockouts and mutational outcomes ordered based on hierarchical clustering with Ward linkage and Euclidean distance. (Signature bar plots) Mutational profiles of the identified signatures, with bars denoting the frequency of each mutational outcome. (Bottom heatmap) Depletion in exposure of each identified signature per gene knockout, as the log2 fold-change between the exposure of the gene knockout sample and the geometric mean exposure of the non-targeting controls. Key DSB repair genes and complexes highlighted.

Clustering of mutational outcomes based on the same mutational frequency changes showed consistent effects across the three target sequence contexts for certain categories of mutations (Fig. 2 central heatmap and MT bar): shorter deletions (*<*4 bp) frequently co-occurred with insertions, while longer deletions with microhomologies correlated with templated insertion (TINS) events. Such co-occurrences suggest that the mutational outcomes could be influenced by similar repair processes and genes, independent of target site-specific effects.

Nevertheless, clustering analysis ignores that genes can be involved in multiple repair pathways, and pathways may share responsibility for producing specific mutational outcomes. To capture such relationships, we analyzed the mutational spectra using the NMF-based SigProfiler framework (see Methods). We identified four distinct and reproducible mutational signatures, CH112A-CH112D, each with unique mutational profiles (Fig. 2 bar plots). Signature CH112A showed the highest proportion of longer deletions with larger MH lengths and the highest frequency of templated insertions (Fig. 2 bar plots and Fig. 3). Signature CH112B showed a larger fraction of small deletions and insertions, while CH112C and CH112D were dominated respectively by HDR events and insertions (Fig. 3). We also observed that outcomes such as HDR events were exclusively linked to one signature (CH112D), whereas other outcomes like insertions appeared in more than one signature (CH112B and CH112D) due to co-occurrence with different mutational outcomes in distinct sets of gene knockout spectra. These results show that NMF has the ability to capture more granular relationships between mutational outcomes possibly associated with multiple mutational processes, which could otherwise be missed using conventional clustering focusing on global similarities across mutational spectra.

**Fig. 3:**
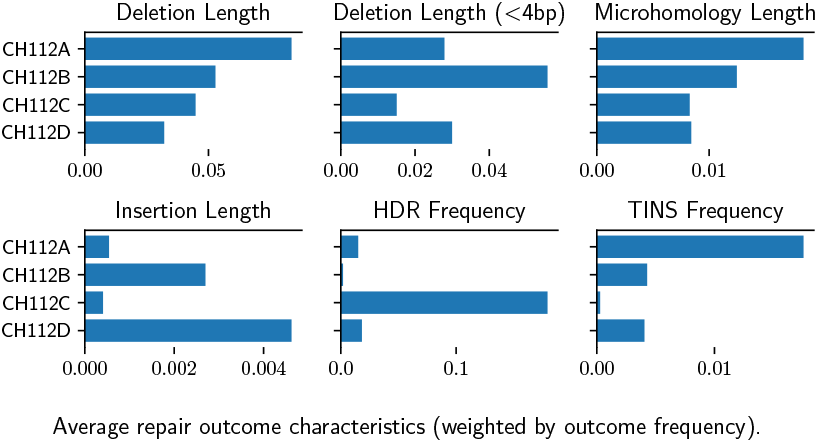
Signature deletion and insertion properties. Per signature average of deletion and insertion outcome properties. Outcome properties: deletion length, small deletion length (*<* 4bp), deletion microhomology (MH) length, insertion length, homology-directed repair (HDR) frequency, and templated insertion (TINS) frequency. The first four length properties are weighted by the signature frequency of the respective outcomes.

We also analyzed the effect of each gene knockout on the contribution of the different signatures to the observed mutational spectrum, by looking at changes in the NMF-estimated signature exposures between gene knockout and controls (see Methods). The diverse impact across genes reflected individual and shared responsibilities for distinct mutational patterns (Fig. 2 bottom heatmap). For example, *Polq* gene knockouts singularly reduced CH112A exposures, while FA core complex gene knockouts lowered CH112C exposures. In contrast, members of the MRN complex influenced multiple signatures (CH112A-B), reflecting their broader role as DNA damage checkpoint genes with an impact on DSB repair pathway choice (Lamarche *et al*., 2010).

Together, this NMF-based analysis of mutational spectra dissected important biological realities that would be missed byconventional techniques, including the multiplicity of roles played by genes and the complexity of shared mutational outcomes arising via alternative repair mechanisms.

### Signature exposures reveal drivers of mutational patterns

We sought to identify mechanisms underlying each mutational signature by examining the biological functions of genes with a larger impact on signature exposures. Controls (Fig. 4, left) showed a prominent contribution from signature CH112A (median 52.9%) linked to deletions with longer MH, followed by CH112C (median 20.3%) associated with HDR events, CH112B (median 14.4%) producing insertions and small deletions, and CH112D (median 12.3%) dominated by insertions. As for the gene knockouts, the largest effects on exposures systematically pointed to a reduction relative to controls (Fig. 4, right), suggesting that those genes tended to promote the mutational signature of interest. To associate signatures with potential biological functions, we performed a functional enrichment analysis of Gene Ontology terms focusing on the genes causing the largest outlying depletion for each signature (see Methods). We found no significantly enriched terms for signature CH112A; however, its association with long MH deletions and the large depletion in exposure caused by *Polq* knockouts suggested a link with the MMEJ pathway (Figs. 3-4). The remaining three signatures, CH112B-D, were enriched for genes involved in DSB-related processes (Fig. 5). Specifically, CH112C was associated with interstrand cross-link repair, a function of the FA core complex. Signatures CH112B and

**Fig. 4:**
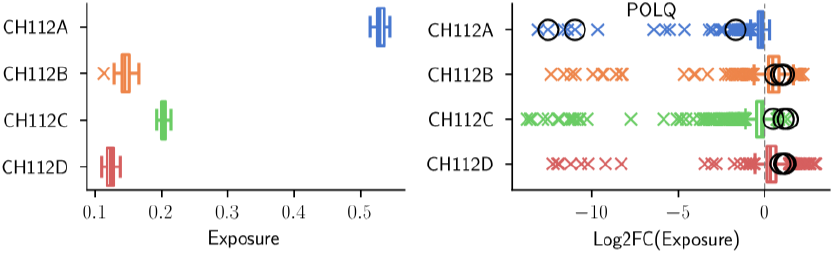
Exposure baseline in controls and change in knockouts. Subfigures: (left) signature exposures of non-targeting controls; (right) impact of every gene knockout on signature exposures, as the log2 fold change between the exposure of the knockout and the geometric mean exposure of non-targeting controls. Black circles highlight *Polq* knockouts. Boxplot: box delimits the interquartile range between 1st and 3rd quartiles (*IQR* = *Q*3 − *Q*1), with a line across the box denoting the median; whiskers indicate the smallest and largest values within [*Q*1 − 1.5 *× IQR, Q*3 + 1.5 *× IQR*], and points beyond them are considered outliers.

**Fig. 5:**
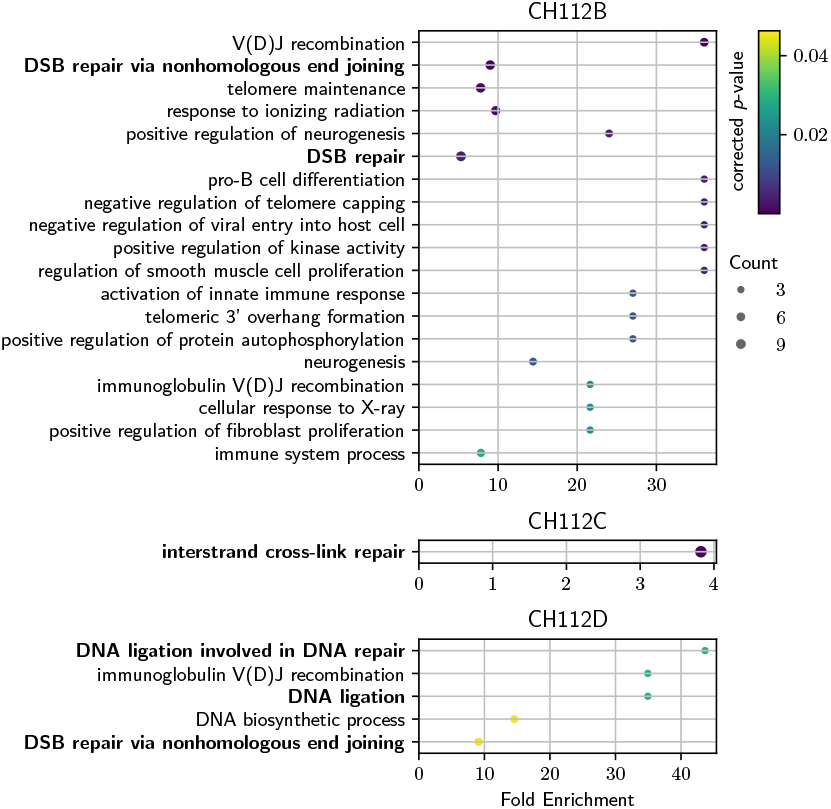
Enriched biological processes for genes involved in signature depletions. Enriched Gene Ontology terms for genes whose knockouts promoted outlying depletion in the exposures of signatures CH112B-D (top to bottom). We analyzed the genes corresponding to outliers below *Q*1 − 1.5 *× IQR* in Fig.4. Signature CH112A did not yield significant results. Each plot shows, for each enriched process (vertical axis): number of genes annotated with the term among the outliers (circle size), fold enrichment as the ratio between the proportions of term-annotated genes among the outliers and among the full set of 766 knocked out genes (horizontal axis) and FDR-corrected p-value (color gradient). Terms directly related to DSB repair are highlighted in boldface.

CH112D were both linked to non-homologous end joining (NHEJ), with CH112D specifically involved in DNA ligation via the ligase IV complex, and a higher frequency of insertions suggesting a role of the ligase IV complex in producing these mutations.

We recovered signatures linked to similar functions from the Barazas2025 screen of 742 genes on one target site (Supplementary Figs. S4-S7), confirming the robustness of the NMF-based analysis. Naturally, the signatures were also similar but not identical across the datasets, as expected given the differences in gene sets, target sites, and sequencing depth. For instance, signature CH112C responsible for HDR events and related to the FA core complex and cross-link repair (Figs. 2/5 seemed to branch into two Barazas2025 signatures (Supplementary Fig. S4): CH32A, producing HDR events and MH deletions, and influenced by FA core complex genes; and CH32E, linked to HDR events and longer MH deletions, influenced by members of the *Trp53bp1* pathway only present in the Barazas2025 dataset (Shieldin complex genes *Shld1* and *Shld2*; CST complex genes *Ctc1, Stn1*, and *Ten1*; and genes *Rif1* and *Mad2l2*, Mirman *et al*. (2018)). Interestingly, the knockout of genes *Rnf8/Rnf168* led to depletion of both signatures CH32A and CH32E, with larger impact on CH32E. This effect was consistent with the role of the *Rnf8/Rnf168* genes as regulators of pathway choice and recruitment of repair factors, including *Trp53bp1* (Nakada, 2016).

Overall, the analysis of exposure profiles enabled us to identify key drivers of mutational signatures, which we further linked to biological functions to obtain fingerprints of DSB repair activity at CRISPR-induced DSBs.

### Exposures suggest DSB repair role for *Dbr1* and *Hnrnpk* genes

We analyzed genes with large knockout signature depletions and no direct annotations with DSB repair terms to infer their potential roles. In both datasets, *Dbr1* (Debranching RNA Lariats 1) emerged as the top unannotated gene. Knockouts of *Dbr1* resulted in a significant depletion of signatures CH112A and CH112C, respectively linked to MMEJ and FA activity. Similarly, knockouts of *Hnrnpk* (Heterogeneous Nuclear Ribonucleoprotein K), present only in the primary dataset, also showed depletion of CH112A and CH112C. It has been suggested that *Hnrnpk* could be a cofactor of p53-mediated DNA damage response, possibly including activation of *Rrm2* as a downstream target involved in nucleotide metabolism (Huarte *et al*., 2010; Wiesmann *et al*., 2017; Braems *et al*., 2022). Our findings based on signature exposures indicated that *Dbr1* and *Hnrnpk* might influence MMEJ and FA, but further experimental validation would be necessary to elucidate their precise roles in DSB repair.

### Exposure analysis challenges existing repair models

Finally, we investigated if the ability of NMF to capture local co-occurrence patterns of subsets of mutational outcomes across subsets of gene knockouts could reveal genes involved in multiple DSB repair pathways. To achieve this, we analyzed the impact of gene knockouts simultaneously across every pair of signatures, focusing our interpretation on genes with known DSB repair pathway associations (Fig. 7). We observed that the knockout of key NHEJ genes and core members of the *Lig4* complex (*Xrcc4* or *Lig4*) promoted a large depletion of signatures CH112B and CH112D, respectively comprising shorter deletions and primarily 1bp insertions. This finding was consistent with the role of the *Lig4* complex in the final ligation step of NHEJ, whose disruption would be expected to hamper the overall NHEJ pathway function. In contrast with the *Lig4* -*Xrcc4* core heterodimer, the accessory factors *Poll* and *Nhej1* did not seem to influence signature CH112B. However, all four genes caused a depletion of signature CH112D (Fig. 6), suggesting they could be responsible for NHEJ-related insertions (Chang *et al*., 2017). Other candidate polymerases typically involved in NHEJ insertions, such as *Polm* (Ramsden, 2011), did not show the same effect on signature CH112D, highlighting the potential of the NMF signature analysis to refine DSB repair function.

**Fig. 6:**
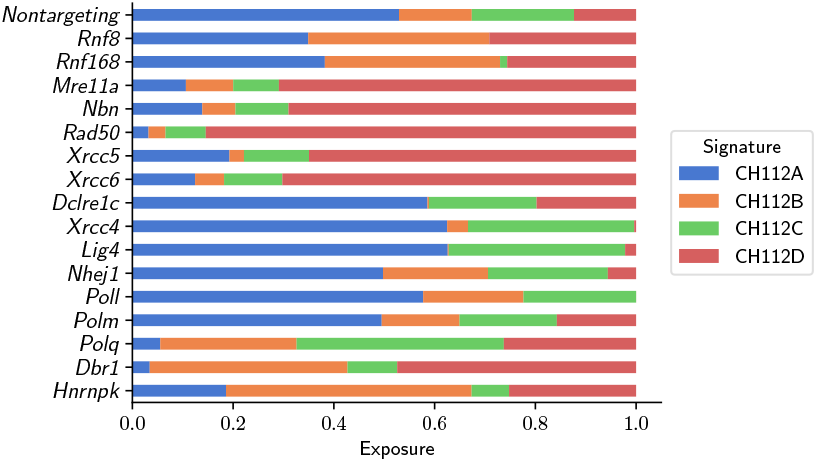
Signature exposure profiles of selected genes. Each signature exposure value was divided by the sum of exposures per knockout and summarized per gene by taking the geometric mean value across knockouts of that gene.

**Fig. 7:**
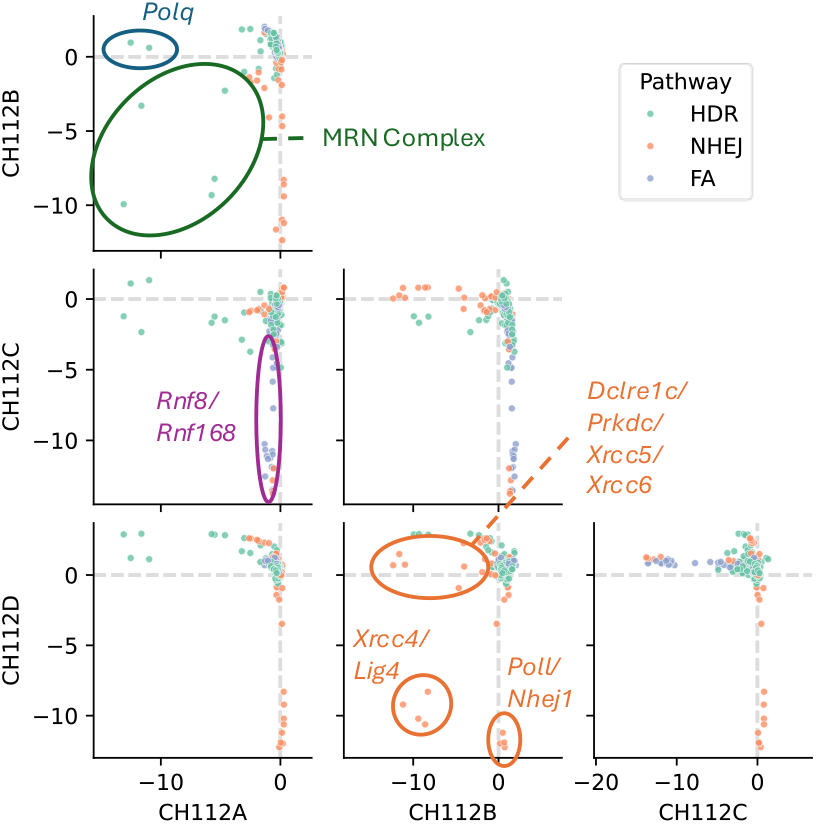
Interplay of signatures based on shared depletion patterns. Pairwise signature depletion scatterplots, with every data point denoting a gene knockout coloured by the DSB repair pathway. The two axes represent signature depletion for the pair of signatures of interest, defined as the log2 fold change in exposure for the gene knockout sample relative to the geometric mean of the non-targeting controls. Grey dashed lines denote zero change. Notable gene knockouts are highlighted using ellipses and labelled.

Interestingly, knockout of other NHEJ genes known as members of the Ku complex and involved in the initiation of repair by NHEJ, *Xrcc5* or *Xrcc6*, led to the depletion of CH112B but not CH112D. Given the association of CH112D with the *Lig4* complex, these results suggested a shift towards Ku-*independent* NHEJ as the dominant repair mechanism in the absence of *Xrcc5* or *Xrcc6*. While classical NHEJ models consider *Xrcc5* /*Xrcc6* /*Dclre1c* required for *Lig4* activity, there is growing evidence for alternative NHEJ circumventing this requirement, though this remains poorly characterized (Schulte-Uentrop *et al*., 2008; Guirouilh-Barbat *et al*., 2007; Fattah *et al*., 2010; Stinson *et al*., 2024).

Knockouts of MRN complex genes reduced CH112A and CH112B activity, indicating the suppression of classical NHEJ and MMEJ pathways. In the absence of MRN complex genes, the Ku-*independent* NHEJ signature CH112D emerged as the predominant repair pathway. Overall, this multi-signature analysis of depletion patterns suggested a high degree of adaptability and cross-talk among DSB repair mechanisms. In particular, elements of how NHEJ compensates for inter- and intra-pathway deficiencies may deserve greater experimental attention.

## Conclusion

In this work, we proposed a computational strategy to infer functions for genes in DSB repair based on high-resolution mutational spectra obtained following gene perturbation screens with CRISPR targeting. Specifically, we identify signatures of mutational activity influenced by both known and candidate DSB repair genes using NMF, attribute these signatures to established DSB repair mechanisms, and link these functions to new genes.

Our NMF analysis of the mutational spectra we generated for a combined set of 307 known and 459 candidate genes revealed an influence of *Dbr1* and *Hnrnpk* knockouts on MMEJ and FA activity, indicating potential roles for these genes in DSB repair. Additionally, signature contributions to mutational spectra revealed two signatures with varying dependencies on members of the *Lig4* complex, representing distinct branches of the NHEJ pathway. This provided evidence of *Lig4* complex activity in the absence of upstream NHEJ factors like *Xrcc5* or *Xrcc6*, suggesting that NHEJ could function independently of *Ku* factors.

Overall, our study highlights the potential of computational approaches for dissecting mutational patterns in CRISPR screens and exploring the genetic landscape of DSB repair. In particular, NMF enables us to identify mutational signatures that serve as fingerprints of repair pathway activity, powering an unsupervised approach to exploring large-scale CRISPR mutational data with the potential for nuanced interpretations and discovery of functional relationships between genes and DSB repair pathways.

## Acknowledgements

The authors acknowledge the High-Performance Compute cluster of the INSY department and the Delft AI cluster at Delft University of Technology. The illustrations in this article were created using BioRender.com.

