## Supplementary Information for "Signatures in CRISPR Mutational Spectra Reveal Role and Interplay of Genes in DNA Repair"

### 1 Supplementary Tables

| Dataset | Target | Replicate | Number of sgRNAs | Number of Frequent Outcomes |
|---|---|---|---|---|
| Primary | T1 | 1 | 2156 | 29 |
| | | 2 | 2235 | 28 |
| | | 3* | **2291** | 28 |
| | T2 | 1 | 2240 | 38 |
| | | 2 | 2249 | 39 |
| | | 3* | **2291** | 40 |
| | T3 | 1 | 2304 | 44 |
| | | 2 | 2291 | 44 |
| | | 3* | **2304** | 44 |
| Barazas2025 | T1 | 1 | 2358 | 32 |
| | | 2* | **2366** | 32 |
| | | 3 | 2350 | 32 |

Table S1: Breakdown of the number of sgRNAs for which mutational spectra were recovered and the number of frequently occurring outcomes per dataset, target site, and replicate. * denotes the replicate that was selected for downstream analysis.

<table>
<thead>
<tr><th></th><th>Type</th><th>Start</th><th>End</th><th>InsSeq</th><th>MHLen</th><th>Repair Product</th></tr>
</thead>
<tbody>
<tr><td>1</td><td>DELETION</td><td>-10.0</td><td>2.0</td><td></td><td>1.0</td><td>CAAGG----------|--AGGGCCTATTTC</td></tr>
<tr><td>2</td><td>DELETION</td><td>-10.0</td><td>5.0</td><td></td><td>3.0</td><td>CAAGG----------|-----GCCTATTTC</td></tr>
<tr><td>3</td><td>DELETION</td><td>-10.0</td><td>6.0</td><td></td><td>2.0</td><td>CAAGG----------|------CCTATTTC</td></tr>
<tr><td>4</td><td>DELETION</td><td>-11.0</td><td>2.0</td><td></td><td>3.0</td><td>CAAG-----------|--AGGGCCTATTTC</td></tr>
<tr><td>5</td><td>DELETION</td><td>-13.0</td><td>17.0</td><td></td><td>3.0</td><td>CA-------------|--------------</td></tr>
<tr><td>6</td><td>DELETION</td><td>-14.0</td><td>8.0</td><td></td><td>2.0</td><td>C--------------|--------TATTTC</td></tr>
<tr><td>7</td><td>DELETION</td><td>-1.0</td><td>5.0</td><td></td><td>3.0</td><td>CAAGGTCGGGCAGG-|-----GCCTATTTC</td></tr>
<tr><td>8</td><td>DELETION</td><td>-1.0</td><td>6.0</td><td></td><td>2.0</td><td>CAAGGTCGGGCAGG-|------CCTATTTC</td></tr>
<tr><td>9</td><td>DELETION</td><td>-24.0</td><td>8.0</td><td></td><td>5.0</td><td>---------------|--------TATTTC</td></tr>
<tr><td>10</td><td>DELETION</td><td>-2.0</td><td>2.0</td><td></td><td>2.0</td><td>CAAGGTCGGGCAG--|--AGGGCCTATTTC</td></tr>
<tr><td>11</td><td>DELETION</td><td>-3.0</td><td>0.0</td><td></td><td>1.0</td><td>CAAGGTCGGGCA---|AGAGGGCCTATTTC</td></tr>
<tr><td>12</td><td>DELETION</td><td>-3.0</td><td>17.0</td><td></td><td>2.0</td><td>CAAGGTCGGGCA---|--------------</td></tr>
<tr><td>13</td><td>DELETION</td><td>-4.0</td><td>7.0</td><td></td><td>4.0</td><td>CAAGGTCGGGC----|-------CTATTTC</td></tr>
<tr><td>14</td><td>DELETION</td><td>-5.0</td><td>5.0</td><td></td><td>2.0</td><td>CAAGGTCGGG-----|-----GCCTATTTC</td></tr>
<tr><td>15</td><td>DELETION</td><td>-6.0</td><td>6.0</td><td></td><td>2.0</td><td>CAAGGTCGG------|------CCTATTTC</td></tr>
<tr><td>16</td><td>DELETION</td><td>-8.0</td><td>7.0</td><td></td><td>1.0</td><td>CAAGGTC--------|-------CTATTTC</td></tr>
<tr><td>17</td><td>DELETION</td><td>-8.0</td><td>8.0</td><td></td><td>1.0</td><td>CAAGGTC--------|--------TATTTC</td></tr>
<tr><td>18</td><td>DELETION</td><td>0.0</td><td>10.0</td><td></td><td>1.0</td><td>CAAGGTCGGGCAGGA|----------TTTC</td></tr>
<tr><td>19</td><td>DELETION</td><td>0.0</td><td>1.0</td><td></td><td>1.0</td><td>CAAGGTCGGGCAGGA|-GAGGGCCTATTTC</td></tr>
<tr><td>20</td><td>DELETION</td><td>0.0</td><td>3.0</td><td></td><td>2.0</td><td>CAAGGTCGGGCAGGA|---GGGCCTATTTC</td></tr>
<tr><td>21</td><td>DELETION</td><td>0.0</td><td>4.0</td><td></td><td>0.0</td><td>CAAGGTCGGGCAGGA|----GGCCTATTTC</td></tr>
<tr><td>22</td><td>DELETION</td><td>0.0</td><td>6.0</td><td></td><td>0.0</td><td>CAAGGTCGGGCAGGA|------CCTATTTC</td></tr>
<tr><td>23</td><td>DELINS</td><td>0.0</td><td>3.0</td><td>G</td><td>0.0</td><td>CAAGGTCGGGCAGGAG---GGGCCTATTTC</td></tr>
<tr><td>24</td><td>HDR</td><td></td><td></td><td>ATTAAGGTACC</td><td></td><td>AGGTCGGGCAGGA<HDR>AGAGGGCCTATT</td></tr>
<tr><td>25</td><td>INSERTION</td><td>0.0</td><td>0.0</td><td>GG</td><td></td><td>AAGGTCGGGCAGGAGGAGAGGGCCTATTTC</td></tr>
<tr><td>26</td><td>INSERTION</td><td>0.0</td><td>0.0</td><td>G</td><td></td><td>CAAGGTCGGGCAGGAGAGAGGGCCTATTTC</td></tr>
<tr><td>27</td><td>INSERTION</td><td>0.0</td><td>0.0</td><td>T</td><td></td><td>CAAGGTCGGGCAGGATAGAGGGCCTATTTC</td></tr>
<tr><td>28</td><td>INSERTION</td><td>1.0</td><td>1.0</td><td>A</td><td></td><td>CAAGGTCGGGCAGGAAAGAGGGCCTATTTC</td></tr>
<tr><td>29</td><td>TINS</td><td></td><td></td><td></td><td></td><td>GGTCGGGCAGGA<TINS>AGAGGGCCTATT</td></tr>
</tbody>
</table>

Table S2: T1 frequently occurring repair outcomes across replicates. Type can be one of DELETION, INSERTION, HDR, or TINS. Start describes the nucleotide position that the deletion begins relative to the cut site, where the cut site itself is position 0. Negative integer values indicate the position is upstream of the cut site. End describes the nucleotide position that the deletion/insertion ends relative to the cut site. InsSeq describes the inserted sequence. MHLen describes the microhomology length for deletions that feature a microhomology, and is 0 otherwise.

<table>
<tr><th></th><th>Type</th><th>Start</th><th>End</th><th>InsSeq</th><th>MHLen</th><th>Repair Product</th></tr>
<tr><td>1</td><td>DELETION</td><td>-1.0</td><td>2.0</td><td></td><td>0.0</td><td>CCTTGGACGCGTAG-|--CCGGTACTAACC</td></tr>
<tr><td>2</td><td>DELETION</td><td>-1.0</td><td>8.0</td><td></td><td>1.0</td><td>CCTTGGACGCGTAG-|--------CTAACC</td></tr>
<tr><td>3</td><td>DELETION</td><td>-2.0</td><td>0.0</td><td></td><td>0.0</td><td>CCTTGGACGCGTA--|GCCCGGTACTAACC</td></tr>
<tr><td>4</td><td>DELETION</td><td>-2.0</td><td>2.0</td><td></td><td>0.0</td><td>CCTTGGACGCGTA--|--CCGGTACTAACC</td></tr>
<tr><td>5</td><td>DELETION</td><td>-2.0</td><td>3.0</td><td></td><td>0.0</td><td>CCTTGGACGCGTA--|---CGGTACTAACC</td></tr>
<tr><td>6</td><td>DELETION</td><td>-3.0</td><td>6.0</td><td></td><td>2.0</td><td>CCTTGGACGCGT---|------TACTAACC</td></tr>
<tr><td>7</td><td>DELETION</td><td>-3.0</td><td>8.0</td><td></td><td>2.0</td><td>CCTTGGACGCGT---|--------CTAACC</td></tr>
<tr><td>8</td><td>DELETION</td><td>-4.0</td><td>11.0</td><td></td><td>2.0</td><td>CCTTGGACGCG----|-----------ACC</td></tr>
<tr><td>9</td><td>DELETION</td><td>-4.0</td><td>1.0</td><td></td><td>2.0</td><td>CCTTGGACGCG----|-CCCGGTACTAACC</td></tr>
<tr><td>10</td><td>DELETION</td><td>-4.0</td><td>2.0</td><td></td><td>2.0</td><td>CCTTGGACGCG----|--CCGGTACTAACC</td></tr>
<tr><td>11</td><td>DELETION</td><td>-5.0</td><td>129.0</td><td></td><td>7.0</td><td>CCTTGGACGC-----|--------------</td></tr>
<tr><td>12</td><td>DELETION</td><td>-5.0</td><td>2.0</td><td></td><td>1.0</td><td>CCTTGGACGC-----|--CCGGTACTAACC</td></tr>
<tr><td>13</td><td>DELETION</td><td>-5.0</td><td>5.0</td><td></td><td>3.0</td><td>CCTTGGACGC-----|-----GTACTAACC</td></tr>
<tr><td>14</td><td>DELETION</td><td>-7.0</td><td>9.0</td><td></td><td>2.0</td><td>CCTTGGAC-------|---------TAACC</td></tr>
<tr><td>15</td><td>DELETION</td><td>-9.0</td><td>0.0</td><td></td><td>0.0</td><td>CCTTGG---------|GCCCGGTACTAACC</td></tr>
<tr><td>16</td><td>DELETION</td><td>-9.0</td><td>1.0</td><td></td><td>0.0</td><td>CCTTGG---------|-CCCGGTACTAACC</td></tr>
<tr><td>17</td><td>DELETION</td><td>-9.0</td><td>4.0</td><td></td><td>2.0</td><td>CCTTGG---------|----GGTACTAACC</td></tr>
<tr><td>18</td><td>DELETION</td><td>0.0</td><td>11.0</td><td></td><td>1.0</td><td>CCTTGGACGCGTAGG|-----------ACC</td></tr>
<tr><td>19</td><td>DELETION</td><td>0.0</td><td>21.0</td><td></td><td>5.0</td><td>CCTTGGACGCGTAGG|--------------</td></tr>
<tr><td>20</td><td>DELETION</td><td>0.0</td><td>2.0</td><td></td><td>1.0</td><td>CCTTGGACGCGTAGG|--CCGGTACTAACC</td></tr>
<tr><td>21</td><td>DELETION</td><td>0.0</td><td>3.0</td><td></td><td>0.0</td><td>CCTTGGACGCGTAGG|---CGGTACTAACC</td></tr>
<tr><td>22</td><td>DELETION</td><td>0.0</td><td>5.0</td><td></td><td>1.0</td><td>CCTTGGACGCGTAGG|-----GTACTAACC</td></tr>
<tr><td>23</td><td>DELETION</td><td>0.0</td><td>7.0</td><td></td><td>2.0</td><td>CCTTGGACGCGTAGG|-------ACTAACC</td></tr>
<tr><td>24</td><td>DELETION</td><td>1.0</td><td>11.0</td><td></td><td>2.0</td><td>CCTTGGACGCGTAGG|G----------ACC</td></tr>
<tr><td>25</td><td>DELETION</td><td>1.0</td><td>2.0</td><td></td><td>1.0</td><td>CCTTGGACGCGTAGG|G-CCGGTACTAACC</td></tr>
<tr><td>26</td><td>DELETION</td><td>1.0</td><td>33.0</td><td></td><td>5.0</td><td>CCTTGGACGCGTAGG|G-------------</td></tr>
<tr><td>27</td><td>DELETION</td><td>1.0</td><td>5.0</td><td></td><td>1.0</td><td>CCTTGGACGCGTAGG|G----GTACTAACC</td></tr>
<tr><td>28</td><td>DELETION</td><td>2.0</td><td>11.0</td><td></td><td>2.0</td><td>CCTTGGACGCGTAGG|GC---------ACC</td></tr>
<tr><td>29</td><td>DELINS</td><td>-1.0</td><td>1.0</td><td>TA</td><td></td><td>CTTGGACGCGTAG-TA-CCCGGTACTAACC</td></tr>
<tr><td>30</td><td>DELINS</td><td>-2.0</td><td>1.0</td><td>TA</td><td></td><td>CTTGGACGCGTA--TA-CCCGGTACTAACC</td></tr>
<tr><td>31</td><td>DELINS</td><td>0.0</td><td>1.0</td><td>GTA</td><td></td><td>CTTGGACGCGTAGGGTA-CCCGGTACTAAC</td></tr>
<tr><td>32</td><td>DELINS</td><td>0.0</td><td>2.0</td><td>G</td><td></td><td>CCTTGGACGCGTAGGG--CCGGTACTAACC</td></tr>
<tr><td>33</td><td>DELINS</td><td>0.0</td><td>2.0</td><td>T</td><td></td><td>CCTTGGACGCGTAGGT--CCGGTACTAACC</td></tr>
<tr><td>34</td><td>HDR</td><td></td><td></td><td>ATTAAGGTACC</td><td></td><td>TTGGACGCGTAGG<HDR>GCCCGGTACTAA</td></tr>
<tr><td>35</td><td>INSERTION</td><td>0.0</td><td>0.0</td><td>A</td><td></td><td>CCTTGGACGCGTAGGAGCCCGGTACTAACC</td></tr>
<tr><td>36</td><td>INSERTION</td><td>0.0</td><td>0.0</td><td>GT</td><td></td><td>CTTGGACGCGTAGGGTGCCCGGTACTAACC</td></tr>
<tr><td>37</td><td>INSERTION</td><td>0.0</td><td>0.0</td><td>G</td><td></td><td>CCTTGGACGCGTAGGGGCCCGGTACTAACC</td></tr>
<tr><td>38</td><td>INSERTION</td><td>0.0</td><td>0.0</td><td>T</td><td></td><td>CCTTGGACGCGTAGGTGCCCGGTACTAACC</td></tr>
<tr><td>39</td><td>INSERTION</td><td>2.0</td><td>2.0</td><td>C</td><td></td><td>CCTTGGACGCGTAGGGCCCCGGTACTAACC</td></tr>
<tr><td>40</td><td>TINS</td><td></td><td></td><td></td><td></td><td>TGGACGCGTAGG<TINS>GCCCGGTACTAA</td></tr>
</table>

Table S3: T2 frequently occurring repair outcomes across replicates. Type can be one of DELETION, INSERTION, HDR, or TINS. Start describes the nucleotide position that the deletion begins relative to the cut site, where the cut site itself is position 0. Negative integer values indicate the position is upstream of the cut site. End describes the nucleotide position that the deletion/insertion ends relative to the cut site. InsSeq describes the inserted sequence. MHLen describes the microhomology length for deletions that feature a microhomology, and is 0 otherwise.

| | Type | Start | End | InsSeq | MHLen | Repair Product |
|---|---|---|---|---|---|---|
| 1 | DELETION | -12.0 | 4.0 | | 2.0 | CCC------------\|----GGCCCGGTAC |
| 2 | DELETION | -13.0 | 9.0 | | 2.0 | CC-------------\|---------GGTAC |
| 3 | DELETION | -16.0 | 11.0 | | 3.0 | ---------------\|-----------TAC |
| 4 | DELETION | -1.0 | 30.0 | | 4.0 | CCCGACCTTGGACG-\|-------------- |
| 5 | DELETION | -1.0 | 7.0 | | 1.0 | CCCGACCTTGGACG-\|-------CCGGTAC |
| 6 | DELETION | -1.0 | 8.0 | | 1.0 | CCCGACCTTGGACG-\|--------CGGTAC |
| 7 | DELETION | -23.0 | 7.0 | | 4.0 | ---------------\|-------CCGGTAC |
| 8 | DELETION | -2.0 | 0.0 | | 0.0 | CCCGACCTTGGAC--\|GTAGGGCCCGGTAC |
| 9 | DELETION | -2.0 | 2.0 | | 0.0 | CCCGACCTTGGAC--\|--AGGGCCCGGTAC |
| 10 | DELETION | -2.0 | 4.0 | | 1.0 | CCCGACCTTGGAC--\|----GGCCCGGTAC |
| 11 | DELETION | -3.0 | 1.0 | | 0.0 | CCCGACCTTGGA---\|-TAGGGCCCGGTAC |
| 12 | DELETION | -3.0 | 2.0 | | 1.0 | CCCGACCTTGGA---\|--AGGGCCCGGTAC |
| 13 | DELETION | -3.0 | 5.0 | | 1.0 | CCCGACCTTGGA---\|-----GCCCGGTAC |
| 14 | DELETION | -3.0 | 6.0 | | 2.0 | CCCGACCTTGGA---\|------CCCGGTAC |
| 15 | DELETION | -3.0 | 8.0 | | 0.0 | CCCGACCTTGGA---\|--------CGGTAC |
| 16 | DELETION | -4.0 | 12.0 | | 1.0 | CCCGACCTTGG----\|------------AC |
| 17 | DELETION | -4.0 | 28.0 | | 5.0 | CCCGACCTTGG----\|-------------- |
| 18 | DELETION | -4.0 | 2.0 | | 1.0 | CCCGACCTTGG----\|--AGGGCCCGGTAC |
| 19 | DELETION | -4.0 | 6.0 | | 2.0 | CCCGACCTTGG----\|------CCCGGTAC |
| 20 | DELETION | -5.0 | 16.0 | | 5.0 | CCCGACCTTG-----\|-------------- |
| 21 | DELETION | -5.0 | 2.0 | | 2.0 | CCCGACCTTG-----\|--AGGGCCCGGTAC |
| 22 | DELETION | -6.0 | 10.0 | | 2.0 | CCCGACCTT------\|----------GTAC |
| 23 | DELETION | -6.0 | 3.0 | | 1.0 | CCCGACCTT------\|---GGGCCCGGTAC |
| 24 | DELETION | -7.0 | 10.0 | | 2.0 | CCCGACCT-------\|----------GTAC |
| 25 | DELETION | -7.0 | 3.0 | | 1.0 | CCCGACCT-------\|---GGGCCCGGTAC |
| 26 | DELETION | -8.0 | 1.0 | | 2.0 | CCCGACC--------\|-TAGGGCCCGGTAC |
| 27 | DELETION | -8.0 | 3.0 | | 2.0 | CCCGACC--------\|---GGGCCCGGTAC |
| 28 | DELETION | -9.0 | 1.0 | | 0.0 | CCCGAC---------\|-TAGGGCCCGGTAC |
| 29 | DELETION | -9.0 | 6.0 | | 2.0 | CCCGAC---------\|------CCCGGTAC |
| 30 | DELETION | 0.0 | 16.0 | | 1.0 | CCCGACCTTGGACGC\|-------------- |
| 31 | DELETION | 0.0 | 1.0 | | 0.0 | CCCGACCTTGGACGC\|-TAGGGCCCGGTAC |
| 32 | DELETION | 0.0 | 5.0 | | 1.0 | CCCGACCTTGGACGC\|-----GCCCGGTAC |
| 33 | DELETION | 0.0 | 6.0 | | 1.0 | CCCGACCTTGGACGC\|------CCCGGTAC |
| 34 | DELETION | 1.0 | 10.0 | | 1.0 | CCCGACCTTGGACGC\|G---------GTAC |
| 35 | DELETION | 1.0 | 13.0 | | 2.0 | CCCGACCTTGGACGC\|G------------C |
| 36 | DELETION | 1.0 | 3.0 | | 2.0 | CCCGACCTTGGACGC\|G--GGGCCCGGTAC |
| 37 | DELETION | 1.0 | 9.0 | | 1.0 | CCCGACCTTGGACGC\|G--------GGTAC |
| 38 | DELETION | 2.0 | 6.0 | | 1.0 | CCCGACCTTGGACGC\|GT----CCCGGTAC |
| 39 | DELINS | -1.0 | 2.0 | G | | CCCGACCTTGGACG-G--AGGGCCCGGTAC |
| 40 | DELINS | -3.0 | 2.0 | G | | CCCGACCTTGGA---G--AGGGCCCGGTAC |
| 41 | DELINS | 0.0 | 2.0 | CGG | | CCGACCTTGGACGCCGG--AGGGCCCGGTA |
| 42 | HDR | | | ATTAAGGTACC | | CGACCTTGGACGC<HDR>GTAGGGCCCGGT |
| 43 | INSERTION | 0.0 | 0.0 | C | | CCCGACCTTGGACGCCGTAGGGCCCGGTAC |
| 44 | INSERTION | 1.0 | 1.0 | G | | CCCGACCTTGGACGCGGTAGGGCCCGGTAC |
| 45 | TINS | | | | | GACCTTGGACGC<TINS>GTAGGGCCCGGT |

Table S4: T3 frequently occurring repair outcomes across replicates. Type can be one of DELETION, INSERTION, HDR, or TINS. Start describes the nucleotide position that the deletion begins relative to the cut site, where the cut site itself is position 0. Negative integer values indicate the position is upstream of the cut site. End describes the nucleotide position that the deletion/insertion ends relative to the cut site. InsSeq describes the inserted sequence. MHLen describes the microhomology length for deletions that feature a microhomology, and is 0 otherwise.

| | Type | Start | End | InsSeq | MHLen | Repair Product |
|---|---|---|---|---|---|---|
| 1 | DELETION | -10.0 | 2.0 | | 1.0 | CAAGG----------\|--AGGGCCTATTTC |
| 2 | DELETION | -10.0 | 5.0 | | 3.0 | CAAGG----------\|-----GCCTATTTC |
| 3 | DELETION | -10.0 | 6.0 | | 2.0 | CAAGG----------\|------CCTATTTC |
| 4 | DELETION | -11.0 | 2.0 | | 3.0 | CAAG-----------\|--AGGGCCTATTTC |
| 5 | DELETION | -13.0 | 17.0 | | 3.0 | CA-------------\|-------------- |
| 6 | DELETION | -14.0 | 8.0 | | 2.0 | C--------------\|--------TATTTC |
| 7 | DELETION | -1.0 | 5.0 | | 3.0 | CAAGGTCGGGCAGG-\|-----GCCTATTTC |
| 8 | DELETION | -1.0 | 6.0 | | 2.0 | CAAGGTCGGGCAGG-\|------CCTATTTC |
| 9 | DELETION | -24.0 | 8.0 | | 5.0 | ---------------\|--------TATTTC |
| 10 | DELETION | -2.0 | 2.0 | | 2.0 | CAAGGTCGGGCAG--\|--AGGGCCTATTTC |
| 11 | DELETION | -3.0 | 0.0 | | 1.0 | CAAGGTCGGGCA---\|AGAGGGCCTATTTC |
| 12 | DELETION | -3.0 | 17.0 | | 2.0 | CAAGGTCGGGCA---\|-------------- |
| 13 | DELETION | -4.0 | 7.0 | | 4.0 | CAAGGTCGGGC----\|-------CTATTTC |
| 14 | DELETION | -5.0 | 5.0 | | 2.0 | CAAGGTCGGG-----\|-----GCCTATTTC |
| 15 | DELETION | -6.0 | 6.0 | | 2.0 | CAAGGTCGG------\|------CCTATTTC |
| 16 | DELETION | -8.0 | 7.0 | | 1.0 | CAAGGTC--------\|-------CTATTTC |
| 17 | DELETION | -8.0 | 8.0 | | 1.0 | CAAGGTC--------\|--------TATTTC |
| 18 | DELETION | -9.0 | 2.0 | | 0.0 | CAAGGT---------\|--AGGGCCTATTTC |
| 19 | DELETION | 0.0 | 10.0 | | 1.0 | CAAGGTCGGGCAGGA\|----------TTTC |
| 20 | DELETION | 0.0 | 1.0 | | 1.0 | CAAGGTCGGGCAGGA\|-GAGGGCCTATTTC |
| 21 | DELETION | 0.0 | 3.0 | | 2.0 | CAAGGTCGGGCAGGA\|---GGGCCTATTTC |
| 22 | DELETION | 0.0 | 4.0 | | 0.0 | CAAGGTCGGGCAGGA\|----GGCCTATTTC |
| 23 | DELETION | 0.0 | 6.0 | | 0.0 | CAAGGTCGGGCAGGA\|------CCTATTTC |
| 24 | DELETION | 0.0 | 7.0 | | 0.0 | CAAGGTCGGGCAGGA\|-------CTATTTC |
| 25 | DELINS | -3.0 | 1.0 | AG | 0.0 | AAGGTCGGGCA---AG-GAGGGCCTATTTC |
| 26 | DELINS | 0.0 | 3.0 | G | | CAAGGTCGGGCAGGAG---GGGCCTATTTC |
| 27 | HDR | | | ATTAAGGTACC | | AGGTCGGGCAGGA<HDR>AGAGGGCCTATT |
| 28 | INSERTION | 0.0 | 0.0 | GG | | AAGGTCGGGCAGGAGGAGAGGGCCTATTTC |
| 29 | INSERTION | 0.0 | 0.0 | G | | CAAGGTCGGGCAGGAGAGAGGGCCTATTTC |
| 30 | INSERTION | 0.0 | 0.0 | T | | CAAGGTCGGGCAGGATAGAGGGCCTATTTC |
| 31 | INSERTION | 1.0 | 1.0 | ATA | | AAGGTCGGGCAGGAAATAGAGGGCCTATTT |
| 32 | INSERTION | 1.0 | 1.0 | A | | CAAGGTCGGGCAGGAAAGAGGGCCTATTTC |
| 33 | TINS | | | | | GGTCGGGCAGGA<TINS>AGAGGGCCTATT |

Table S5: Barazas2025 frequently occurring repair outcomes across replicates. Type can be one of DELETION, INSERTION, HDR, or TINS. Start describes the nucleotide position that the deletion begins relative to the cut site, where the cut site itself is position 0. Negative integer values indicate the position is upstream of the cut site. End describes the nucleotide position that the deletion/insertion ends relative to the cut site. InsSeq describes the inserted sequence. MHLen describes the microhomology length for deletions that feature a microhomology, and is 0 otherwise.

| Dataset | Target | Replicate | 1 | 2 | 3 |
|---|---|---|---|---|---|
| Primary | T1 | 1 | 1.000 | 0.982 | 0.983 |
| | | 2 | 0.982 | 1.000 | 0.984 |
| | | 3 | 0.983 | 0.984 | 1.000 |
| | T2 | 1 | 1.000 | 0.981 | 0.983 |
| | | 2 | 0.981 | 1.000 | 0.982 |
| | | 3 | 0.982 | 0.983 | 1.000 |
| | T3 | 1 | 1.000 | 0.990 | 0.990 |
| | | 2 | 0.990 | 1.000 | 0.990 |
| | | 3 | 0.990 | 0.990 | 1.000 |
| Barazas2025 | T1 | 1 | 1.000 | 0.993 | 0.992 |
| | | 2 | 0.993 | 1.000 | 0.992 |
| | | 3 | 0.992 | 0.992 | 1.000 |

Table S6: Mean Pearson's correlation coefficient across common genes and outcomes between mutational spectra of replicates for the same target site.

#### 2 Supplementary Figures

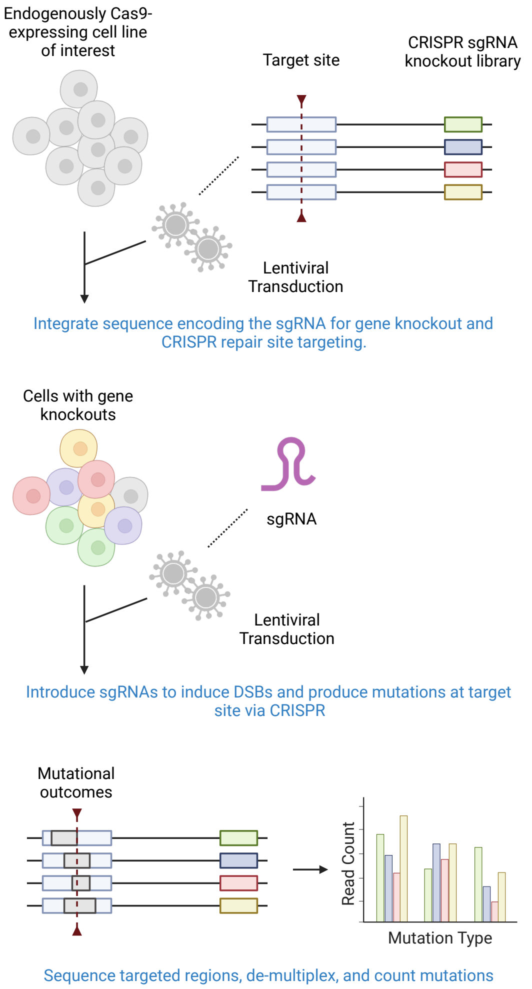

Figure S1: Illustration of CRISPR gene knockout screens with mutational spectra readout. First, sequences are integrated into the genomes of cells via lentiviral transduction. Each sequence contains two elements: *(i)* a sgRNA-encoding region to knockout a single gene, and *(ii)* a region common to all integrated sequences to be targeted with CRISPR to produce the mutational spectra. After genomic integration, several days of cell culture are allowed for genes to be knocked out. Following this, lentiviral transduction is used to introduce sgRNAs targeting the common region to the Cas9-expressing cells. After allowing time for cell culture for DNA cleavage and repair, DNA sequencing was performed to capture the final CRISPR repair products.

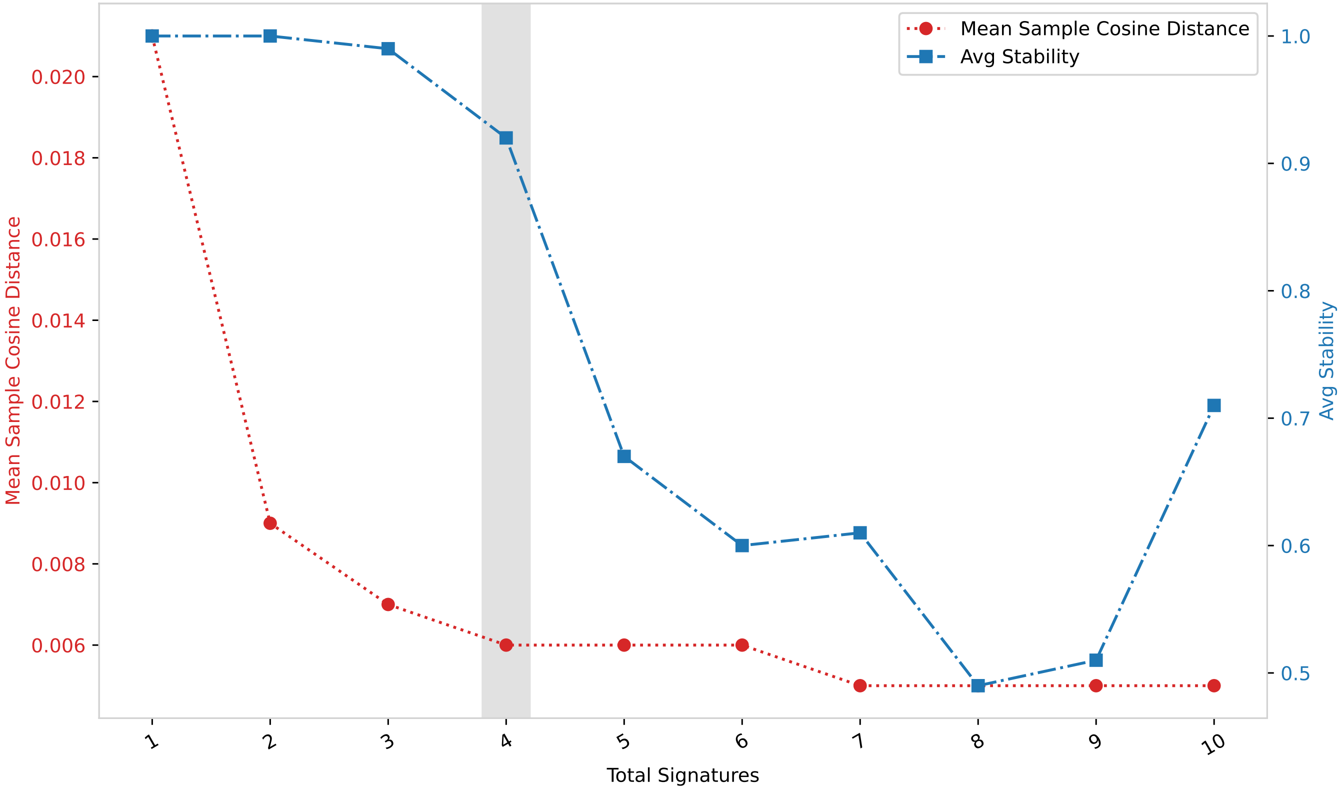

Figure S2: Comparison of NMF solutions for different k values for primary dataset. The left axis represents the reconstruction error as "Mean Sample Cosine Distance". Lower error means better reconstruction. The right axis represents the stability of the solution as the "average silhouette similarity". A higher stability means the solution does not vary a lot between successive runs of the SigProfiler algorithm. The horizontal axis represents the different values tested under for $k$.

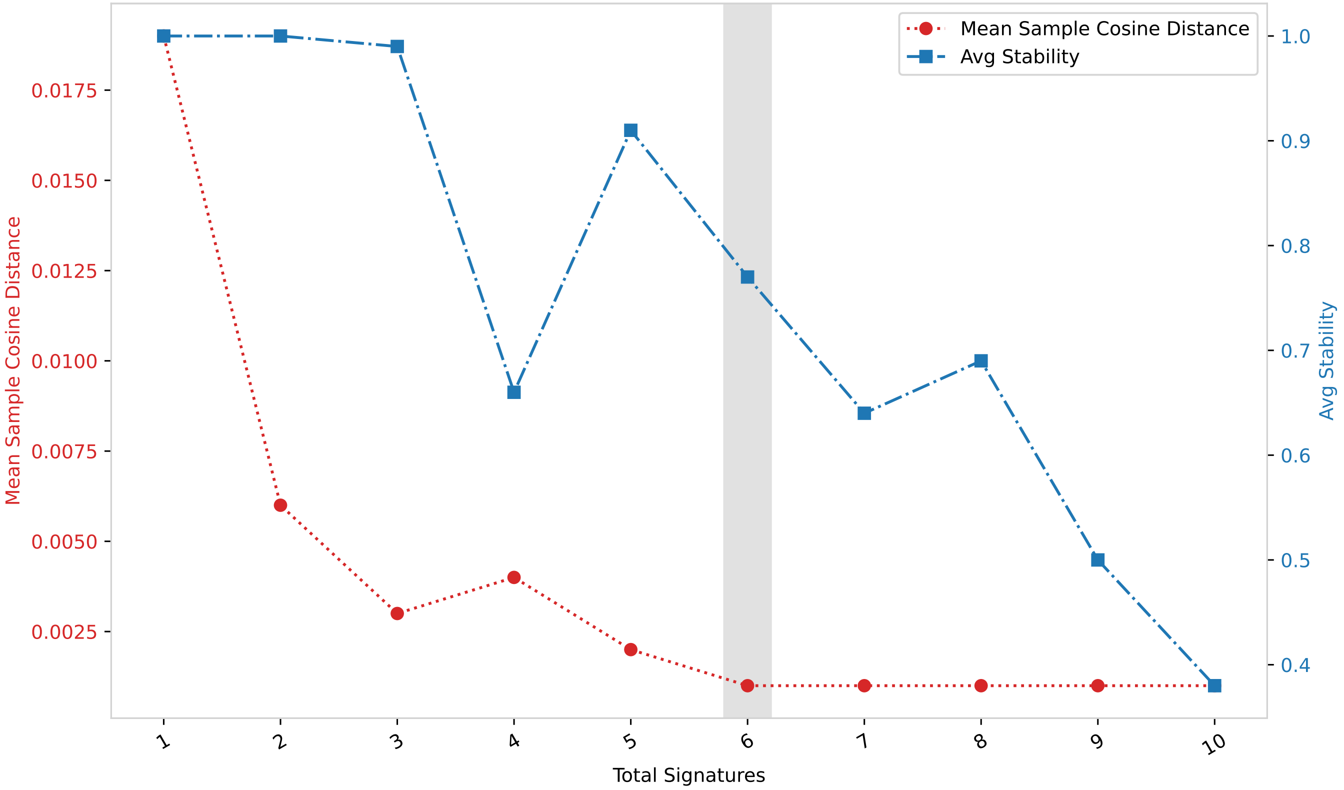

Figure S3: Comparison of NMF solutions for different k values for Barazas2025 dataset. The left axis represents the reconstruction error as ”Mean Sample Cosine Distance”. Lower error means better reconstruction. The right axis represents the stability of the solution as the ”average silhouette similarity”. A higher stability means the solution does not vary a lot between successive runs of the SigProfiler algorithm. The horizontal axis represents the different values tested under for $k$.

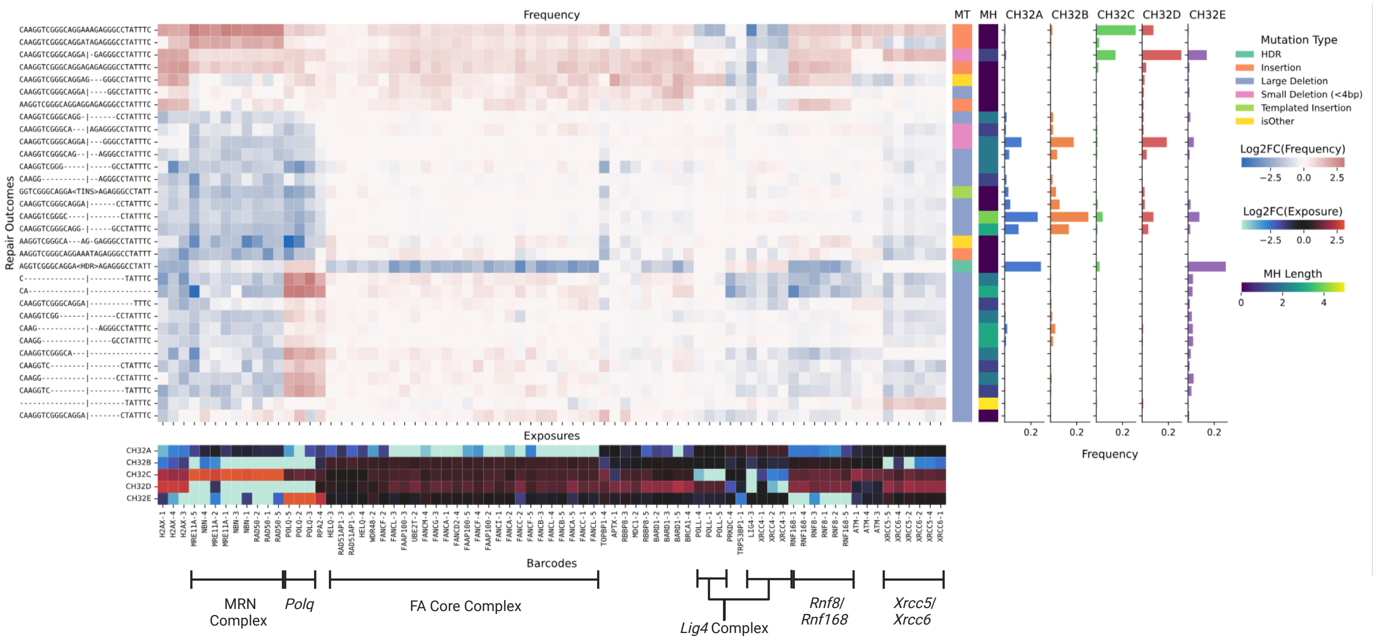

Figure S4: Impact of gene knockouts on mutational spectra and identified mutational signatures for the Barazas2025 dataset. (Central heatmap) Effect of the 75 gene knockouts with the largest outcome redistribution (columns) on every mutational outcome across three target sites (rows), with outcomes further characterized by mutation type (MT) and microhomology length (MH). Effect quantified using the log2 fold-change in outcome frequency between the gene knockout spectrum and the geometric mean of the non-targeting control spectra. Gene knockouts and mutational outcomes ordered based on hierarchical clustering with Ward linkage and Euclidean distance. (Signature bar plots) Mutational profiles of the identified signatures, with bars denoting the frequency of each mutational outcome. (Bottom heatmap) Depletion in exposure of each identified signature per gene knockout, as the log2 fold-change between the exposure of the gene knockout sample and the geometric mean exposure of the non-targeting controls. Key DSB repair genes and complexes highlighted.

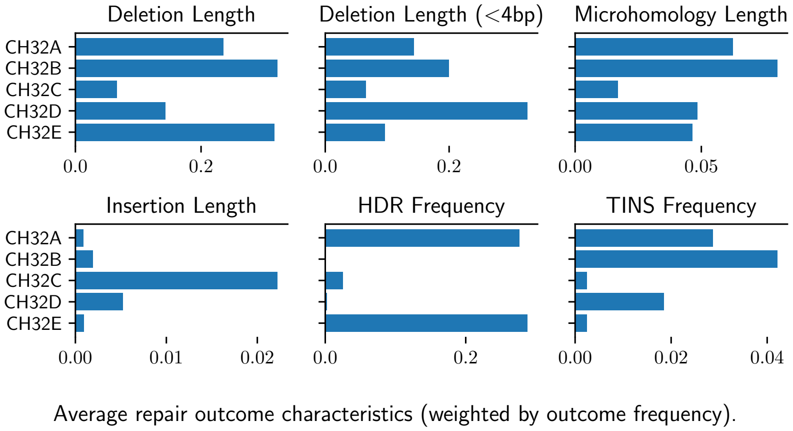

Figure S5: Barazas2025 signature deletion and insertion properties. Per signature average of deletion and insertion outcome properties. Outcome properties: deletion length, small deletion length ($< 4$bp), deletion microhomology (MH) length, insertion length, homology-directed repair (HDR) frequency, and templated insertion (TINS) frequency. The first four length properties are weighted by the signature frequency of the respective outcomes.

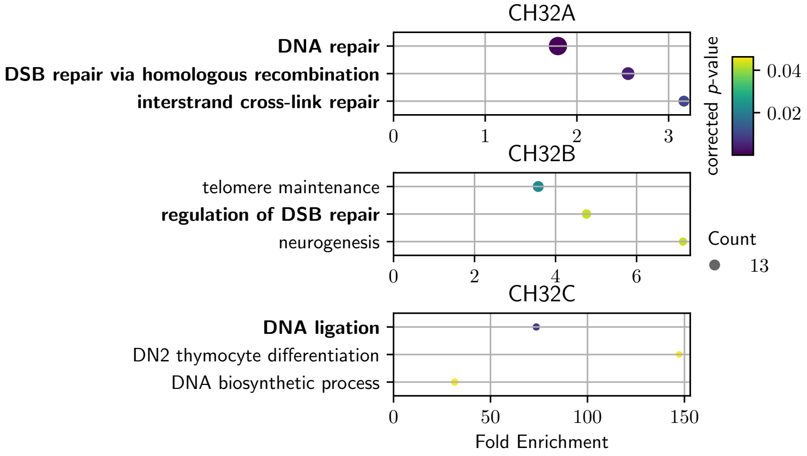

Figure S6: Enriched biological processes for genes involved in Barazas2025 signature depletions. Enriched Gene Ontology terms for genes whose knockouts promoted outlying depletion in the exposures of signatures CH112B-D (top to bottom). We analyzed the genes corresponding to outliers below $Q1 - 1.5 \times IQR$ in Fig.**??**. Signature CH112A did not yield significant results. Each plot shows, for each enriched process (vertical axis): number of genes annotated with the term among the outliers (circle size), fold enrichment as the ratio between the proportions of term-annotated genes among the outliers and among the full set of 766 knocked out genes (horizontal axis) and FDR-corrected p-value (color gradient). Terms directly related to DSB repair are highlighted in boldface.

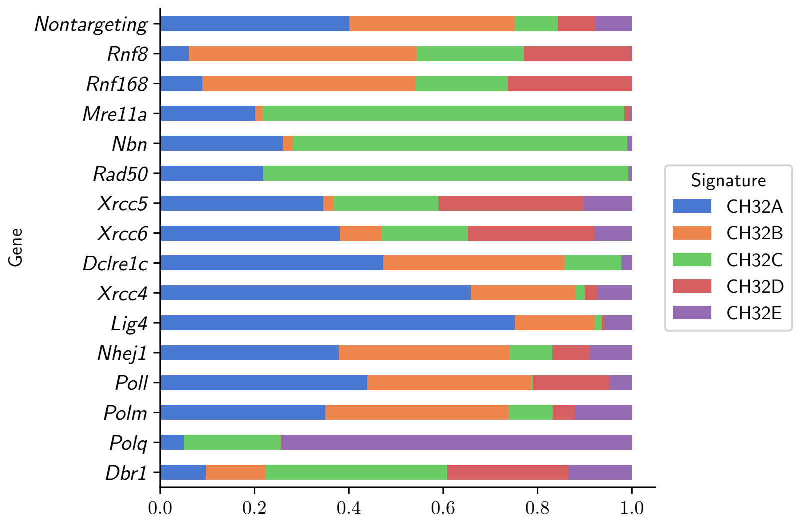

Figure S7: Barazas2025 signature exposure profiles of selected genes. Each signature exposure value was divided by the sum of exposures per knockout, and summarized per gene by taking the geometric mean value. across knockouts of that gene.
